# Specialization for cell-free or cell-to-cell spread of BAC-cloned HCMV strains is determined by factors beyond the UL128-131 and RL13 loci

**DOI:** 10.1101/760611

**Authors:** Eric P. Schultz, Jean-Marc Lanchy, Le Zhang Day, Qin Yu, Christopher Peterson, Jessica Preece, Brent J. Ryckman

## Abstract

It is widely held that clinical isolates of human cytomegalovirus (HCMV) are highly cell-associated, and mutations affecting the UL128-131 and RL13 loci that arise in culture lead to the appearance of a cell-free spread phenotype. The BAC-clone Merlin (ME), expresses abundant UL128-131, is RL13-impaired and produces low infectivity virions in fibroblasts, whereas TB40/e (TB) and TR are low in UL128-131, RL13-intact and produce virions of much higher infectivity. Despite these differences, quantification of spread by flow cytometry revealed remarkably similar spread efficiencies in fibroblasts. In epithelial cells, ME spread more efficiently, consistent with robust UL128-131 expression. Strikingly, ME spread far better than TB or TR in the presence of neutralizing antibodies on both cell types, indicating that ME is not simply deficient at cell-free spread, but is particularly efficient at cell-to-cell spread, whereas TB and TR cell-to-cell spread is poor. Sonically disrupted ME-infected cells contained scant infectivity, suggesting that the efficient cell-to-cell spread mechanism of ME depends on features of the intact cells such as junctions or intracellular trafficking processes. Even when UL128-131 was transcriptional repressed, cell-to-cell spread of ME was still more efficient than TB or TR. Moreover, RL13 expression comparably reduced both cell-free and cell-to-cell spread of all three strains, suggesting that it acts at a stage of assembly and/or egress common to both routes of spread. Thus, HCMV strains can be highly specialized for either for cell-free or cell-to-cell spread and these phenotypes are determined by factors beyond the UL128-131 or RL13 loci.

**IMPORTANCE:** Both cell-free and cell-to-cell spread are likely important for the natural biology of HCMV. In culture, strains clearly differ in their capacity for cell-free spread as a result of differences in the quantity and infectivity of extracellular released progeny. However, it has been unclear whether “cell-associated” phenotypes are simply the result of poor cell-free spread, or are indicative of particularly efficient cell-to-cell spread mechanisms. By measuring the kinetics of spread at early time points, we were able to show that HCMV strains can be highly specialized to either cell-free or cell-to-cell mechanisms, and this was not strictly linked the efficiency of cell-free spread. Our results provide a conceptual approach to evaluating intervention strategies for their ability to limit cell-free or cell-to-cell spread as independent processes.

## INTRODUCTION

Intervention approaches to control human cytomegalovirus (HCMV) infection, including DNA replication inhibitors such as ganciclovir, and vaccines designed to elicit neutralizing antibodies have had limited success (1, 2), and this may be due in part to the complex genetic diversity of HCMV circulating within human populations (3–9). Basic annotations of the 235kbp HCMV genome identify 165 canonical open reading frames (ORFs), although there is evidence of extensive transcription and translation beyond these loci (10, 11). While nucleotide polymorphisms are found throughout the genome, most sequences are well conserved. 21 of the 165 canonical ORFs show high nucleotide diversity and are distributed as islands throughout the genome. Several groups have reported evidence of frequent recombination within the more conserved regions, which may essentially mix and match the more diverse loci into many different genotype combinations (3, 4, 7, 8). Adding to this complexity, some studies have shown evidence of gene inactivating mutations (pseudogenes), and gene deletions (7, 8). While some infected individuals may harbor relatively pure populations of HCMV genotypes, complex, multiple genotype infections, and sequential infections by genetically distinct populations also occur. Cloning of HCMV from clinical samples on bacterial artificial chromosomes (BAC) has resulted in numerous, genetically distinct strains for use in laboratory studies. It is not clear how the genetic and phenotypic differences among BAC-cloned HCMVs might reflect their natural diversity.

The glycoprotein (g) H/L complex is functionally conserved among the herpesvirus family and is involved in the initial receptor engagement and the regulation of the membrane fusion protein gB (reviewed in (12)). The HCMV gH/gL is present in the virion envelope as a gH/gL/gO complex (13, 14), a gH/gL/UL128-131 complex (15–18), and in complex with gB (19). Transient expression of HCMV gH/gL and gB is sufficient to drive cell-cell fusion, but is unclear which forms of gH/gL contribute to membrane fusion during HCMV infection (20). The gH/gL/gO and gH/gL/UL128-131 complexes bind to various cell surface receptors through the gO and UL128-131 domains (21–23) and these interactions are important for entry into a broad range of cell types. gH/gL/gO is likely critical for infection of most, or all cell types, with fibroblasts, epithelial, and endothelial cells being the most extensively studied (24–27). Mutants of HCMV lacking gO are deficient at adsorption to cells and soluble gH/gL/gO blocks such adsorption (25, 28). In contrast, gH/gL/UL128-131 is dispensable for infection of fibroblasts and neuronal cells but critical for epithelial and endothelial cells, monocyte-macrophages, and dendritic cells (16, 18, 29–31). Soluble gH/gL/UL128-131 did not block virus adsorption, but there is evidence that engagement of receptors can elicit signal transduction pathways that influence the nature of the resulting infection pathway (21, 28, 30). Murine and guinea pig CMVs contain homologous gH/gL complexes that play similar roles in cell tropism, although there are some differences in the reported requirements for the complexes for infection of different cell types (32–37).

We, and others have noted striking phenotypic variation among the HCMV strains TB40/e (TB), TR, and Merlin (ME) in terms of content of the gH/gL complexes in the virion envelope and the corollary effects on entry and tropism. TB and TR contain gH/gL mostly in the form of gH/gL/gO, and very little gH/gL/UL128-131, whereas ME virions contain overall lower amounts of gH/gL and this is mostly in the form of gH/gL/UL128-131 (27). The genetic polymorphism(s) responsible for these differences are unclear. Murrell et al. described a G>T mutation in the UL128 locus of TB that when engineered into ME, reduced the assembly of gH/gL/UL128-131 through effects on mRNA splicing (38). However, this did not fully explain the observed strain differences, as TR is also low in gH/gL/UL128-131 but is congenic to ME at this locus (27). Zhang et al. showed that expression of gO during replication is lower in ME-infected cells as compared to TR, but again the genetic correlates of this difference were not clearly identified (39). The cell-free infectivity of HCMV strains on both fibroblasts and epithelial cells correlates with the amounts of gH/gL/gO in the virion envelope; TB is by far the most infectious, followed by TR, while ME virions are poorly infectious (27). Repression of the UL131 promoter in ME resulted in virions with dramatically reduced amounts of gH/gL/UL128-131, somewhat higher levels of gH/gL/gO, and dramatically improved cell-free infectivity (27, 40).

HCMV can spread through monolayer cell cultures by diffusion of cell-free virus in the culture supernatant or by a more direct cell-to-cell mode, but the mechanistic distinctions between these types of spread are not well characterized. Moreover, the pathway of cell-to-cell spread for HCMV has not been extensively studied. There have been suggestions of limited fusion between infected and uninfected cells allowing the transfer of subviral components, but the efficiency of these processes to facilitate the spread of HCMV infection is not clear (45–47). Syncytium formation has been observed in HCMV infected cultures, but it is not clear whether these cell-cell fusions contribute to viral spread, or merely represent the coalescence of late-stage infected cells, long after progeny virus has exited and spread to adjacent cells (48, 49). Different HCMV strains can be characterized as cell-free or cell-associated based on the appearance of the foci formed in a culture monolayer. For example, ME is considered cell-associated due to formation of tightly localized foci whereas TB forms more diffuse focal pattern characteristic of cell-free spread (40, 41). However, it is not clear whether the apparent cell-associated nature of ME simply reflects the lack of efficient cell-free spread, or a specific mechanism to facilitate efficient cell-to-cell spread. Nor is it clear how strains like ME and TB compare in their ability to spread cell-to-cell. Finally, the genetic loci responsible for the apparent cell-free and cell-associated natures of strains are not clear. Some reports have monitored foci size in the presence of neutralizing antibodies and attributed the cell-associated nature of ME to the expression of the UL128-131 and RL13 loci (24, 42). It seems clear that either gH/gL/gO or gH/gL/UL128-131 is required for cell-to-cell spread on fibroblasts, whereas gH/gL/UL128-131 seems to be required for cell-to-cell spread in epithelial or endothelial cells (24, 26, 43, 44). Here, we report the use of flow cytometry to compare the spread of HCMV TB, TR, and ME in fibroblasts and epithelial cells at early time points. The results showed that these strains differ not only in their ability to spread cell-free, but also cell-to-cell and that neither UL128-131 nor the RL13 are the determinate loci for the mode of spread.

## RESULTS

### Representative HCMV strains differ not only in their efficiencies of cell-free spread, but also in their ability to spread cell-to-cell

Focal sizes and patterns can vary considerably among strains of HCMV and likely reflect differences in their proclivity for cell-free versus cell-to-cell modes of spread. Representative focal spread patterns of GFP-expressing HCMV strains TB, TR, and ME are depicted in Figure 1. Cultures of fibroblasts and epithelial cells were infected at low multiplicities and spread was monitored by fluorescence microscopy over 18 days. In fibroblasts, the spread of all three strains was localized to small, tight foci for the first 6 days. By day 12, TB and TR began to show signs of more diffuse spread that continued to increase through day 18, whereas ME foci remained generally smaller and more localized over the entire experiment (Fig 1A). In contrast, foci in epithelial cell cultures for all three strains were smaller and more tightly localized throughout the experiment (Fig. 1B). The more diffuse focal patterns of TB and TR in fibroblasts were likely indicative of efficient cell-free spread, whereas the localized spread of ME was suggestive of less efficient cell-free spread. This was consistent with our previous observations that progeny virus released to culture supernatants by TB and TR were far more infectious than extracellular ME progeny (27). While similar focal spread characterizations have been reported by other groups (40) (24) (38) (50), such comparisons of focal size and patterns are not easily quantified and do not clearly distinguish whether smaller, more localized foci form as a passive result of inefficient cell-free spread, or rather reflect an active preference for a specialized cell-to-cell mechanism. Moreover, while it is clear that these strains of HCMV can differ in their ability to spread via cell-free progeny, whether they also differ in their ability to spread cell-to-cell has not been addressed.

**Figure 1.**
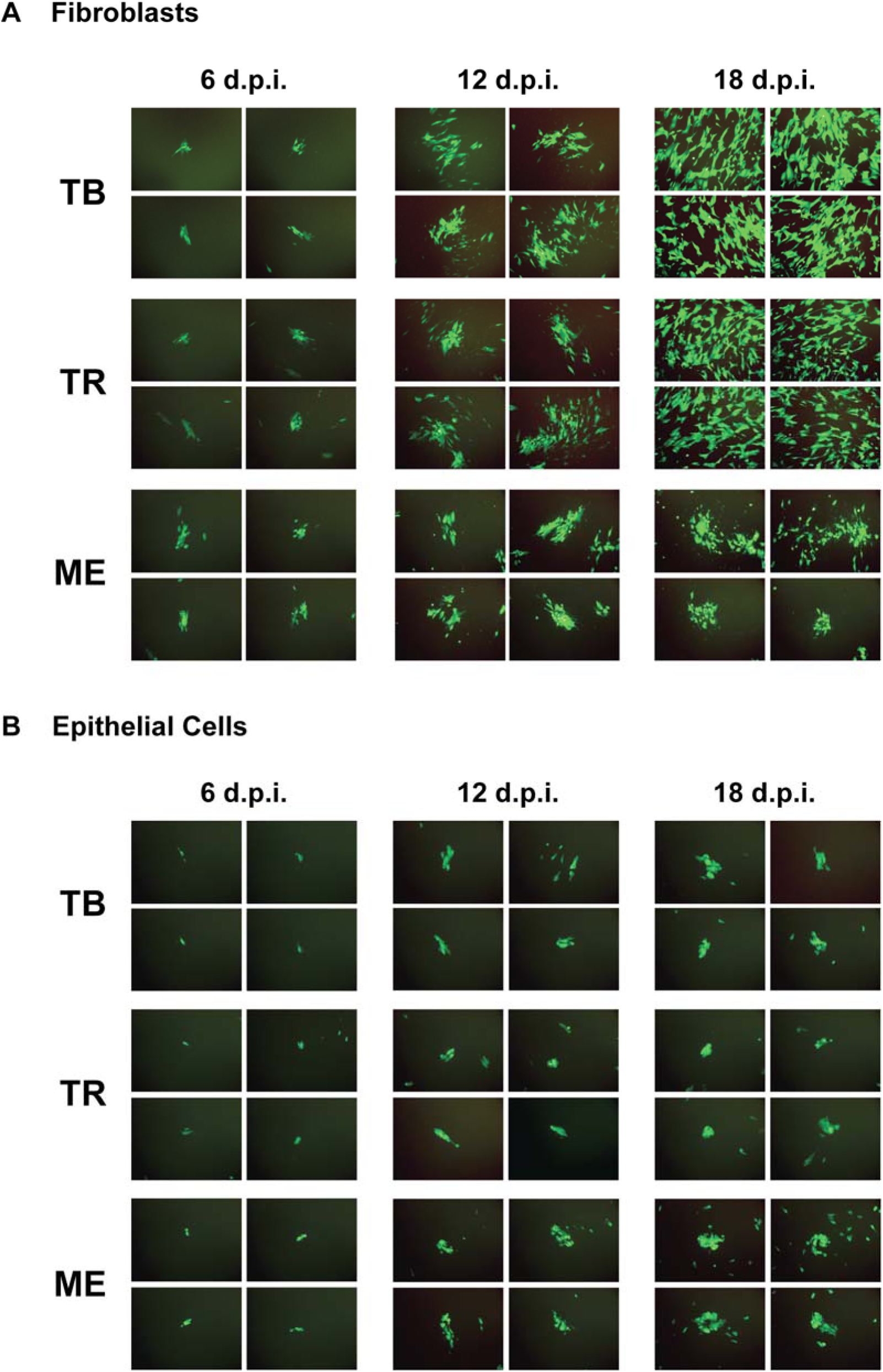
Comparison of focal spread patterns among distinct HCMV strains. Confluent monolayers of fibroblasts (A) or epithelial cells (B) were infected at MOI 0.001 with GFP-expressing HCMV TB, TR, or ME. Foci were documented by fluorescence microscopy at 6, 12, and 18 days post infection. Four representative fields (10X) are shown for each.

For quantitative comparisons of spread among HCMV strains, experiments similar to those described for Figure 1 were performed, but instead of microscopy, flow cytometry was used to measure the increasing number of infected (GFP-expressing) cells over the first 12 days of replication. In fibroblasts, the number of infected cells for both TB and TR increased exponentially over 12 days, and fit well to a log-linear regression, indicating a constant rate of spread over the course of the experiment (Fig 2A, B). In contrast, ME spread considerably faster between days 3 and 6, and then spread slower between days 9 and 12. The average spread rates (LN GFP+ cells/day as indicated by the regression slopes in Fig. 2B) of all three strains were within 1.2-fold of each other, with only the difference between TB and TR reaching statistical significance (Fig. 2C). This was somewhat unexpected, as ME has been described to replicate poorly compared to other strains (38). Thus, the growth of the smaller, more tightly localized foci of ME were characterized by initially rapid expansion followed by progressive slowing, whereas the more diffuse foci of TB and TR were characterized by a constant rate of expansion. These results highlight how visual analyses of focal size and pattern fail to detect the rapid, early focal expansion of ME. In epithelial cells, spread rates for all three stains were lower than in fibroblasts, and there were greater differences between strains. ME was approximately 2-fold faster than TR and 1.5 fold faster than TB, and all differences were statistically significant (Fig. 2D-F). The more efficient spread by ME on epithelial cells was consistent with the higher levels of gH/gL/UL128-131 as compared to TB and TR (38) (51) (27), but the tightly localized focal patterns suggested inefficient cell-free spread by all three strains on this cell type.

**Figure 2.**
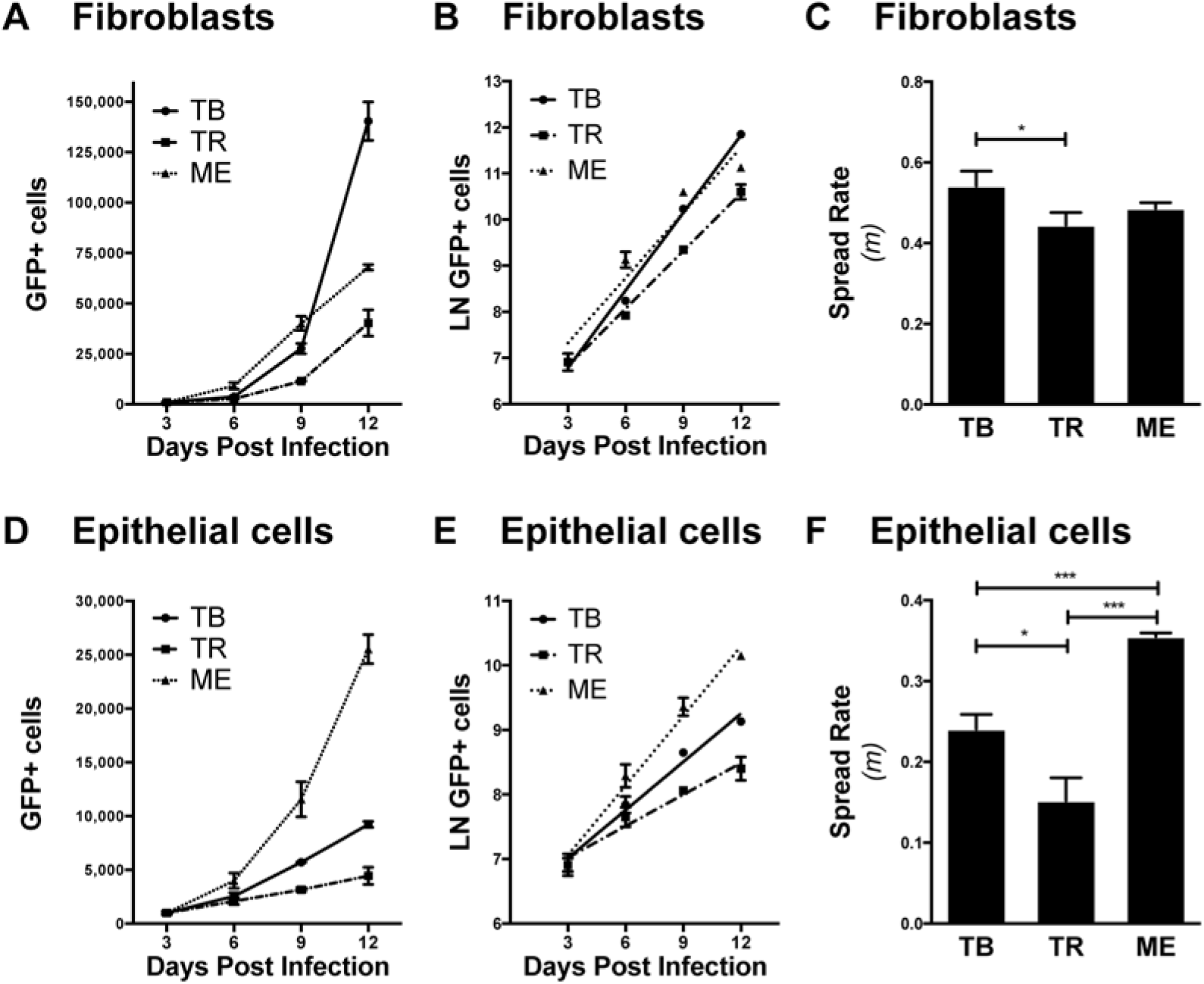
Quantitation of HCMV spread by in fibroblasts and epithelial cells. Confluent 9.5cm^2^ monolayers of fibroblasts (A) or epithelial cells (D) were infected at MOI 0.001 with GFP-expressing HCMV strains TB, TR, or ME. The number of infected cells at 3, 6, 9, and 12 days post infection was determined by flow cytometry. (B and E) LN GFP+ cells is plotted at each time point shown in panels A and D, respectively. (C and F) The average spread rate (LN GFP+ cells per day) as the slope (m) of the regression lines in panels B and E, respectively. All experiments were performed at least three times and error bars represent standard deviation between all experiments (omitted if smaller than the data marker), and p-values were generated using unpaired, two-tailed t-tests (*<0.05, **<0.01, ***<0.001).

Neutralizing antibodies were used to distinguish the contributions of cell-free and cell-to-cell mechanisms to the rate of spread for each strain. Antibodies chosen for these experiments were mAb 14-4b, an anti-gH antibody that likely targets a discontinuous epitope at the membrane proximal region (52, 53), as well as a mixture of rabbit anti-peptide sera that target the epithelial tropism factors UL130 and UL131 (17). The relative potency of these antibodies to neutralize cell-free TB, TR, and ME was verified in neutralization experiments shown in Figure 3. Note that on fibroblasts, 14-4b was approximately 10-fold more potent against ME than for TB and TR, and there was a residual 20% of TR infectivity that was resistant even at very high antibody concentrations (Fig 3A). On epithelial cells, potency of neutralization by anti-UL130/131 antibodies was more similar among the strains, and complete neutralization of each was achieved (Fig. 3B).

**Figure 3.**
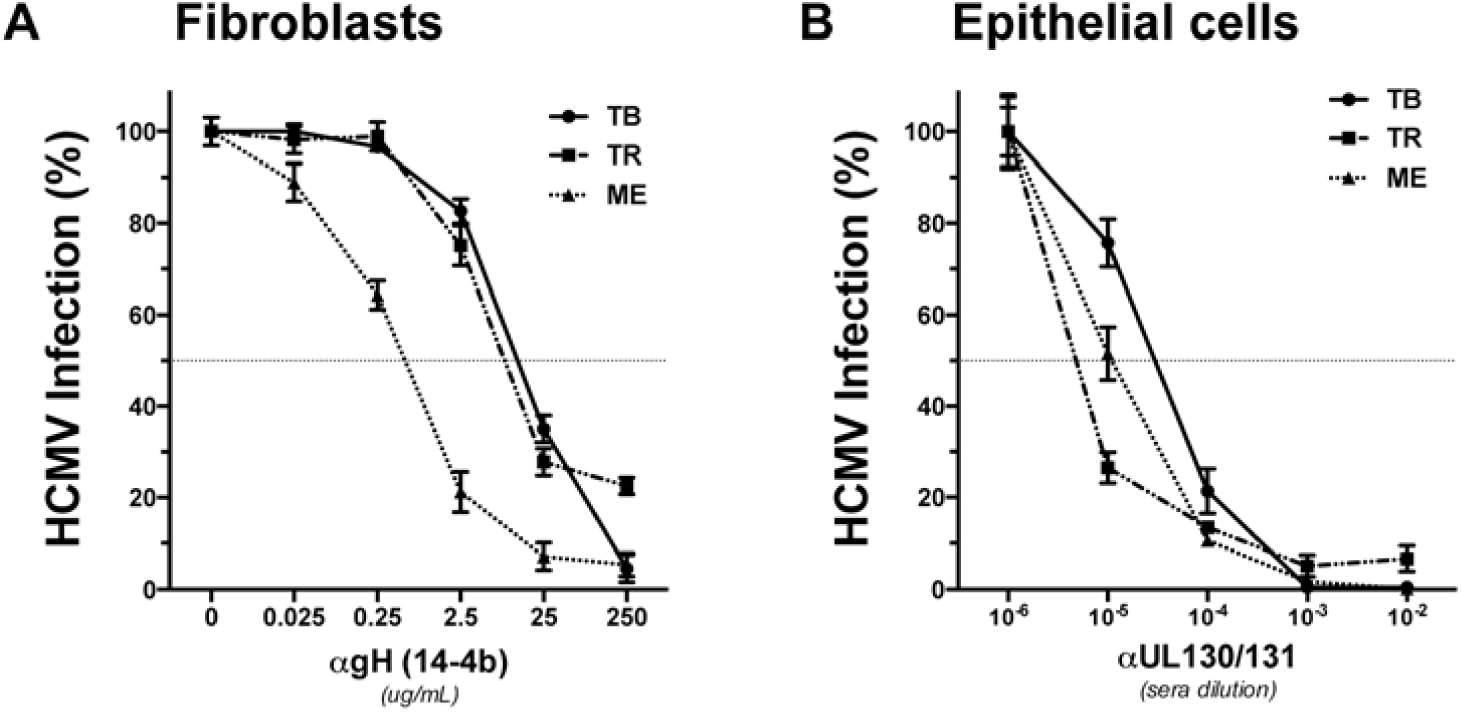
Antibody neutralization of cell-free HCMV. Equal numbers (genomes/mL) of fibroblast-derived (A) or epithelial-derived (B) HCMV TB, TR, or ME virions were incubated with multiple concentrations of anti-gH mAb 14-4b (A) or anti-UL130/131 rabbit sera (B) for 1hr at RT. Remaining infectivity was determined by titration on the matched producer cell type, and plotted as percent of no antibody control. All experiments were performed in triplicate and error bars represent standard deviation.

Spread rates of each strain in either fibroblast or epithelial cell cultures were then determined in the presence or absence of anti-gH mAb 14-4b or anti-UL130/131 sera at concentrations sufficient for maximal neutralization of cell-free virions. In fibroblast cultures, anti-gH 14-4b reduced the spread rate of TB and TR by 70% and 55%, respectively, whereas ME spread was reduced by only 25% (Fig. 4 A-C). The apparent resistance of ME spread was especially noteworthy since cell-free ME was more sensitive to neutralization than either TB or TR (Fig. 3A). In all cases, the apparent antibody resistant spread was greater than the spread in the presence of ganciclovir, even for TR, which is known to harbor resistance mutations in the UL97 kinase (54). This demonstrated that the apparent antibody resistant spread was not simply the transfer of cytoplasmic contents from infected to uninfected cells, but rather represented *bona fide* viral spread dependent on the production of new viral genomes. These results supported the hypothesis that spread of TB in fibroblasts predominately involved cell-free virions that were highly sensitive to neutralizing antibodies, whereas ME spread predominately by a distinct, antibody resistant cell-to-cell mechanism. The intermediate antibody sensitivity of TR spread might indicate intermediate contributions of cell-free and cell-to-cell spread mechanisms. However, it may also reflect the lack of complete neutralization of cell-free TR as described in Fig 3A. In epithelial cell cultures, the presence of anti-UL130/131 antibodies resulted in 20-30% reductions in spread rates for all three strains, indicating predominantly cell-to-cell modes of spread in this cell type (Fig 4 D-F). Again, ME spread was most efficient in the presence of neutralizing antibodies, despite TB and TR displaying similar resistance over 12 days. While virions derived from fibroblasts are well characterized (51) (27), less is known about virions produced in epithelial cells. To this end, virion quantity and infectivity generated from epithelial cells infected with TB, TR, or ME was determined using subcellular fractionation, and compared to fibroblasts.

**Figure 4.**
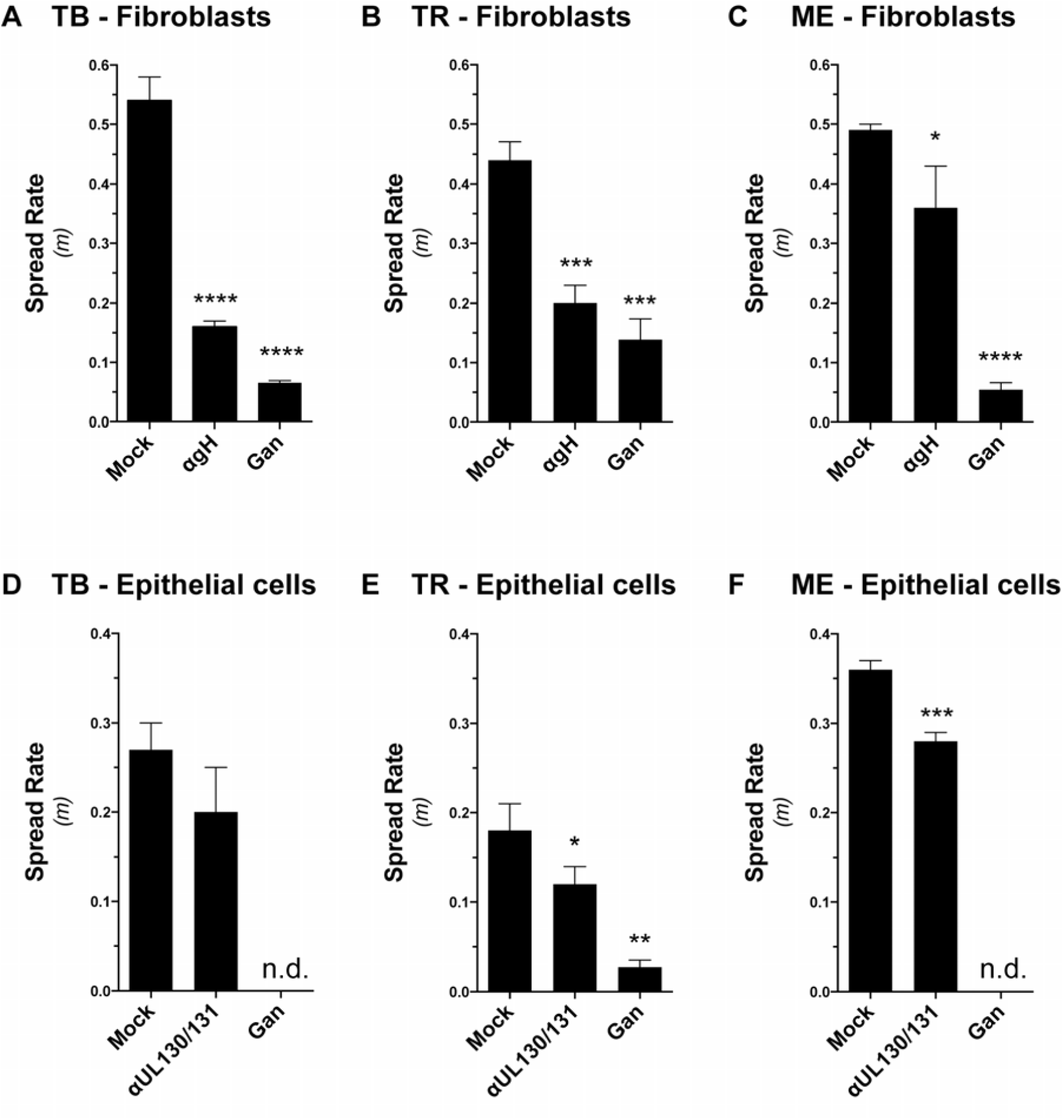
Effects of neutralizing antibodies on spread of HCMV strains in fibroblasts and epithelial cells. Confluent 9.5cm^2^ monolayers of fibroblasts or epithelial cells were infected at MOI 0.001 with GFP-expressing HCMV strains TB, TR, or ME and allowed to spread in the presence of either anti-gH mAb 14-4b (A-C), anti-UL130/131 rabbit sera (D-F), or ganciclovir (all panels). The average spread rates over 12 days (LN GFP+ cells per day; as in Fig 2) for TB, TR, and ME on fibroblasts (A-C) and epithelial cells (D-F) are shown. Conditions where no spread was detected are designated “n.d.”. All experiments were performed three times and error bars represent standard deviation. P-values were generated by unpaired, two-tailed t-test vs. mock (*<0.05, **<0.01, ***<0.001).

### HCMV ME is uniquely efficient at cell-to-cell spread via a mechanism that depends on the intact cell monolayer, and overcomes the low infectivity of progeny virus

The efficient cell-free spread of TB in fibroblasts was likely a function of the quantity and infectivity of progeny virus release to culture supernatants. Similarly, it was expected that highly efficient cell-to-cell spread of ME would correlate with the quantity and/or infectivity of progeny virus within the infected cells, or “cell-associated virus”. To test these hypotheses, infected cultures were separated into supernatants and cells, and the cells further fractionated into nuclear, and cytoplasmic portions. This fractionation allowed for more detailed comparisons of DNA replication and nuclear egress between strains. The quantity and infectivity of progeny virus within the infected cells and in the supernatants were then determined, and results shown in Figure 5.

**Fig 5.**
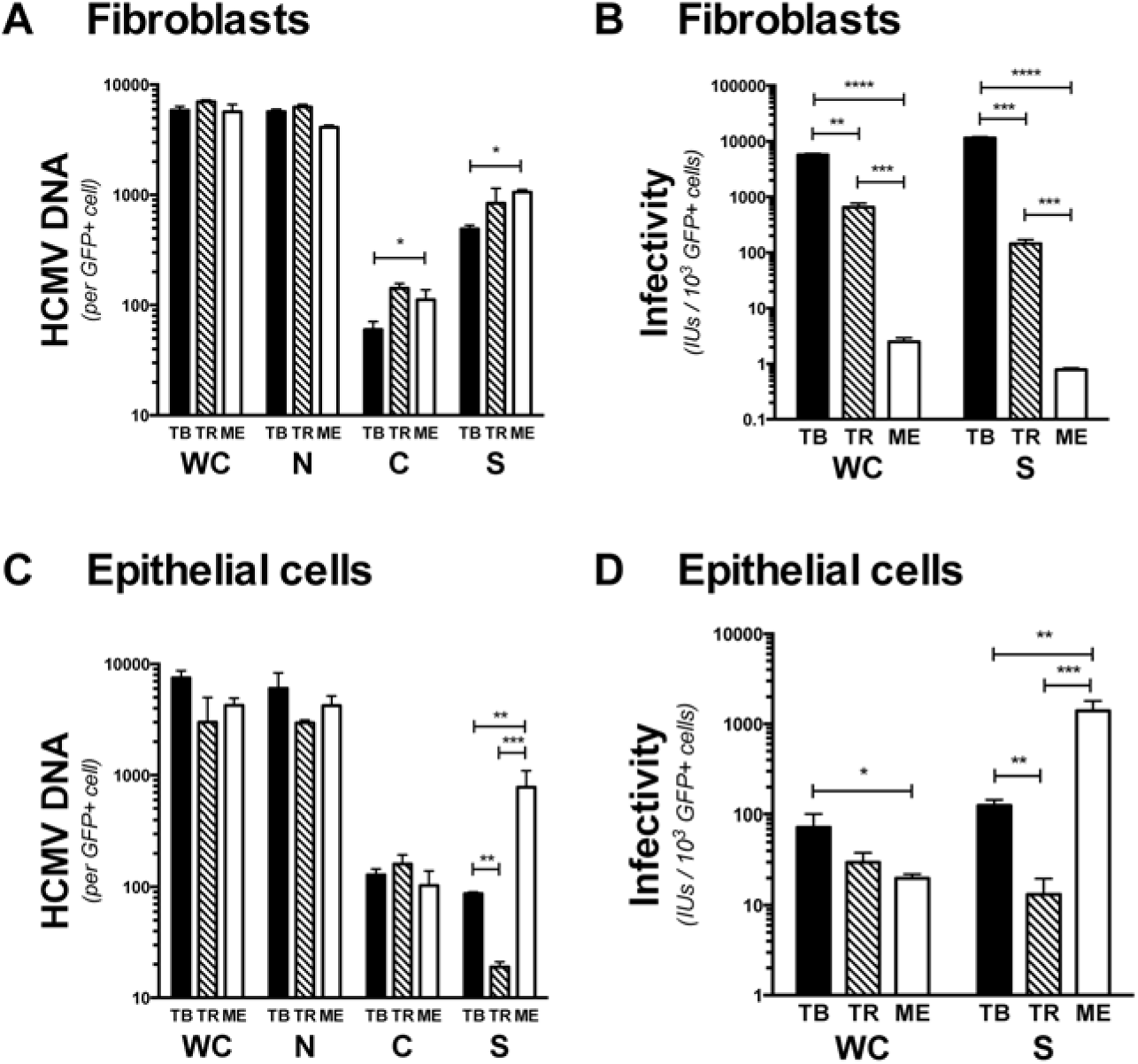
Production and infectivity of cell-free and cell-associated HCMV progeny virus. Replicate cultures of fibroblasts or epithelial cells were infected at MOI 1 with TB, TR, or ME. 7 days later infected cells were quantified by flow cytometry for GFP expression. *(A and C).* Cultures were fractionated into whole cell (WC), nuclear (N), cytoplasmic (C), and supernatant (S), and viral genomes were determined by qPCR. The number of HCMV genomes in each fraction is plotted per GFP+ cell. *(B and D).* Infectious units were determined from whole cell sonicates (WC) and culture supernatants (S) by titration on the matched producer cell type. Total infectious units are plotted per 1000 GFP+ cells. *All panels.* Error bars represent the standard deviation of three separate experiments, and p-values were generated using unpaired, two-tailed t-tests (*<0.05, **<0.01, ***<0.001, ****<0.0001).

In fibroblasts, total intracellular viral genome accumulation was comparable for all three strains (Fig 5A). The vast majority was in the nuclear fraction, likely in the form of non-infectious, unpackaged genomes and non-tegumented, non-enveloped nucleocapsids. The number of viral genomes in cytoplasmic and supernatant fractions was slightly lower for TB-infected fibroblasts than for either TR or ME. At least a portion of these cytoplasmic and supernatant genomes would be expected to represent the infectious progeny that could contribute to either cell-to-cell or cell-free spread, respectively. Despite the lower numbers of progeny genomes for TB in cytoplasmic and supernatant fractions, these corresponded to log-folds greater cell-associated (i.e., sonically disrupted cells) and cell-free (i.e., supernatant) infectivity per cell as compared to TR or ME (Fig. 5B). The greater infectivity of TB, despite lower virion numbers was in agreement with our previous measures of lower specific infectivity ratios (genome/PFU) for TB compared to TR and ME (27), and was positively correlated with the relative efficiencies of cell-free spread for these strains. Surprisingly, cell-to-cell spread was inversely correlated to the levels of cell-associated infectivity, with cell-to-cell spread of ME being highly efficient despite producing the lowest detectable cell-associated infectivity.

In epithelial cells, accumulation of whole cell, cytoplasmic, and nuclear progeny genomes were similar among all three strains (Fig. 5C). In contrast, ME released nearly 10-fold more progeny genomes to culture supernatants than did TB, and TR was by far the least efficient at releasing progeny to supernatants. These results suggest major differences among strains in the later assembly and release stages of the replication cycle in epithelial cells. As was observed in fibroblasts, ME had the lowest cell-associated infectivity in epithelial cells, despite the more efficient cell-to-cell spread in this cell type (Fig 5D and 4F). Infectivity in epithelial cell culture supernatants mirrored the quantities of genomes, indicating that the epithelial-derived virions of all three strains are of comparable specific infectivity. This was in stark contrast to the drastic differences in specific infectivity among these strains when produced in fibroblasts (Fig 5 A and B) (27).

Taken together, these results confirmed that the robust cell-free spread of TB and TR in fibroblasts was associated with the release of highly infectious progeny virus to the culture supernatant and that the poor cell-free spread of ME was due to the low infectivity of the cell-free progeny. Surprisingly, ME was much more efficient at cell-to-cell spread than TB despite having far less intracellular infectious progeny than TB or TR. In epithelial cell cultures, the cell-free characteristics for all of the strains were more similar to one another than they were on fibroblasts, and this was consistent with a comparably small contribution of cell-free spread for all three strains in this cell type. Thus, spread by all three strains in epithelial cells was predominantly cell-to-cell, and ME was by far the most efficient whereas TR was notably inefficient. The cell-to-cell spread of ME thus depends on efficient trafficking of progeny to adjacent cells, and this seems to overcome the poor infectivity of the intracellular progeny.

### The highly efficient cell-to-cell spread mechanism of HCMV ME is not determined by the high expression of UL128-131

The cell-associated nature of ME has been linked to the high expression levels of UL128-131 and the corollary poor infectivity of cell-free ME virions (24, 27, 40, 42). To address this in our quantitative spread assay, we made use of the previously described tetracycline (Tet) repression system developed by Stanton et al. (40). Briefly, the BAC clone of ME used in these studies contained tetracycline-operator (TetO) sequences in the promoter of UL131. We previously showed that replication of this recombinant ME in HFFFtet fibroblasts, which express the tetracycline-repressor (TetR) protein, produced extracellular virus with dramatically reduced gH/gL/UL128-131, slightly higher gH/gL/gO and as a result, greatly improved cell-free infectivity (27). We refer to ME propagated under conditions of UL131 repression as “Merlin-T” (MT).

Direct comparisons of ME spread rates in nHDF the HFFFtet cells were complicated by the fact that in addition to expressing TetR, HFFFtet were also telomerase immortalized and this could have independent effects on the HCMV replication cycle. Moreover, since we also wished to address the role of UL128-131 expression on spread in epithelial cells, two new TetR-expressing cell lines were generated; nHDFtet and ARPEtet. A luciferase reporter assay demonstrated efficient repression of TetO-containing promoters in each of these cell lines (Fig 6A). Consistent with our previous reports, ME virions produced from nHDF contained high levels of gH/gL/UL128-131 and low amounts of gH/gL/gO, whereas MT virions produced in HFFFtet contained far less gH/gL/UL128-131 but more gH/gL/gO (Fig 6B) (27). MT produced in the new nHDFtet cell line showed a similar reduction of gH/gL/UL128-131 and increase in gH/gL/gO, and MT virions derived from either tetR-expressing fibroblasts cell line were dramatically more infectious than ME derived from nHDF cells (Figs 6 B and C). ME virions produced in control ARPE cells were gH/gL/UL128-131-rich and, as in fibroblasts, Tet-repression dramatically reduced amounts of gH/gL/UL128-131 in epithelial-derived MT virions (Fig 6B). However, unlike fibroblast-derived MT, epithelial-derived MT did not contain increased gH/gL/gO and the infectivity was actually reduced (Figs 6B and C).

**Fig 6.**
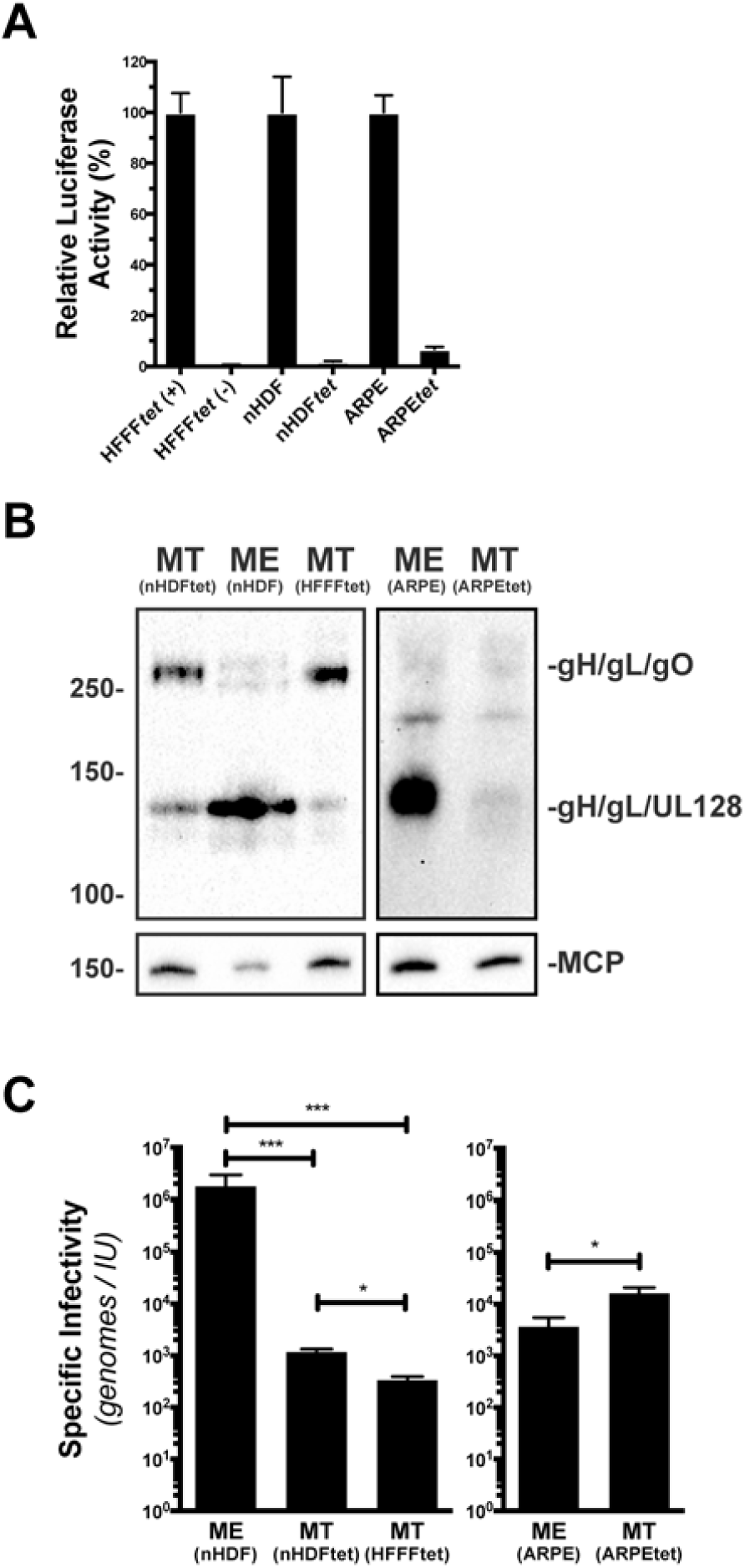
Repression of gH/gL/UL128-131 expression from HCMV ME in fibroblasts and epithelial cells. (A) Tetracycline-repressor protein (tetR) expressing fibroblasts (HFFFtet, nHDFtet) or epithelial cells (ARPEtet) were transfected with TetO-firefly luciferase reporter and the relative luciferase activity was measured. HFFFtet were treated with doxycycline (+) or left untreated (-). (B) Non-reducing (top) or reducing (bottom) immunoblot analyses of MT or ME virions derived from TetR-expressing or control cells, respectively. Blots were probed with anti-gL antibodies (top) or anti-MCP antibodies (bottom). (C) Specific infectivity (genomes/IU) of MT and ME virions derived from culture supernatants of TetR-expressing or control cells, respectively. Titrations of fibroblasts-derived virus were performed of nHDFs, epithelial-derived virus on ARPEs. All experiments were performed in triplicate and error bars represent standard deviation. P-values were generated by unpaired, two-tailed t-test (*<0.05, **<0.01, ***<0.001).

In nHDF fibroblasts, MT spread slightly faster than ME, consistent with a gain in the ability to spread cell-free as a result of the improved virion infectivity (Fig 7A). However, in the presence of neutralizing antibody 14-4b, the spread of ME and MT was indistinguishable. Similar experiments were performed with HFFFtet cells, which generate MT. The high-gH/gL/UL128-131 condition of ME was generated by addition of doxycycline to the cultures to block the TetR repression of UL131 (Fig 6A). Spread rates were generally lower in HFFFtet cells compared to nHDF cells, but as expected, MT spread more efficiently than ME, even in the presence of neutralizing antibodies (Figs 7 A and B). Thus, in both fibroblast systems, the efficiency of ME cell-to-cell spread was not dependent on the high levels of gH/gL/UL128-131. Nor was the apparent cell-to-cell spread a default mechanism of poor cell-free spread, and these results demonstrate a previously unappreciated distinction of the mechanisms of cell-free and cell-to-cell spread.

**Figure 7.**
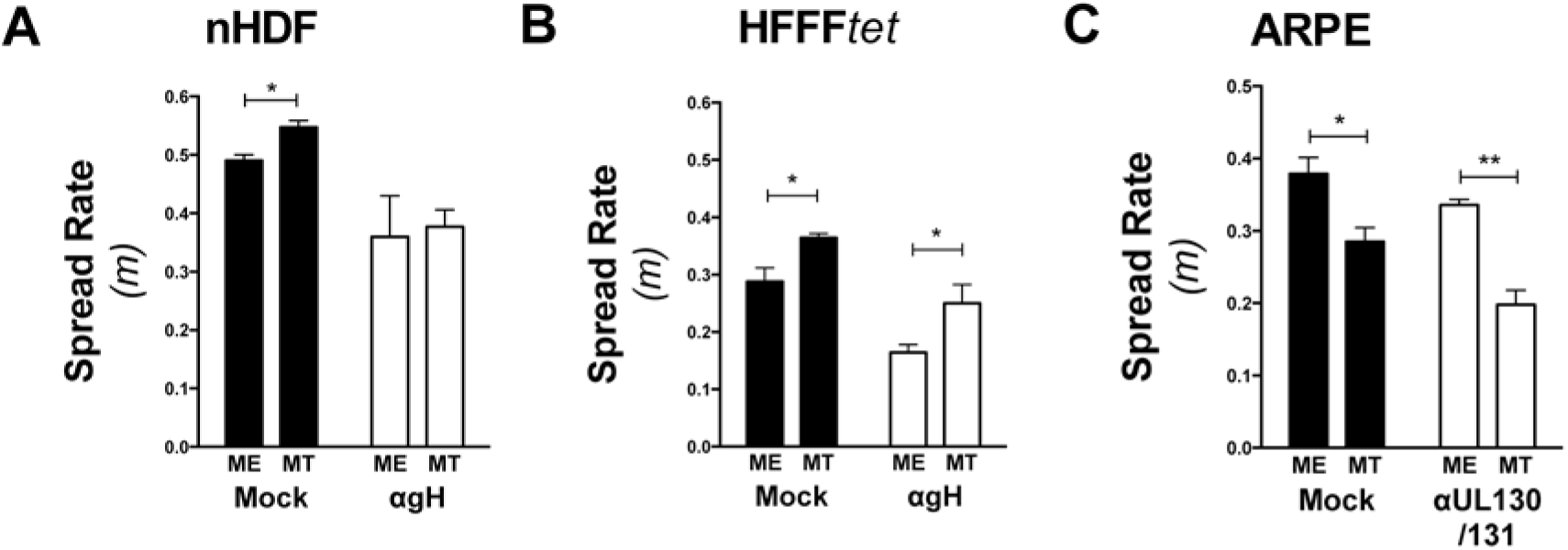
Effect of gH/gL/UL128-131 repression on HCMV ME spread in fibroblasts and epithelial cells. Confluent 9.5cm^2^ monolayers of (A) nHDF or nHDFtet, for ME and MT, respectively, (B) HFFFtet (+) or (-) doxycycline, for ME and MT, respectively, or (C) ARPE or ARPEtet, for ME and MT, respectively were infected with at MOI 0.001 with GFP-expressing HCMV ME/MT. Spread rates in the presence of either anti-gH mAb 14-4b (A and B), or anti-UL130/131 rabbit sera (C) were determined as described in Figure 2. Plotted are averages and standard deviations for three independent experiments. P-values were generated by unpaired, two-tailed t-test (*<0.05, **<0.01).

In epithelial cells, repression of gH/gL/UL128-131 resulted in a decrease in average spread rate (from 0.38 to 0.29), and small increase in the sensitivity of that spread to neutralizing anti-UL130/131 antibodies (from 10% to 30% inhibition) (Fig 7B). The reduction in spread rate was consistent with the important role of gH/gL/UL128-131 for HCMV in these cells. One interpretation of the small increase in antibody sensitivity was a shift towards increased cell-free spread. However, this seemed unlikely since epithelial-derived MT were actually less infectious than epithelial-derived ME (Fig 6C). Thus, it seems more likely that the observed spread of MT was still predominantly cell-to-cell, but that the reduced gH/gL/UL128-131 allowed for more efficient blocking of the cell-to-cell spread by these antibodies. It was notable that the spread rates of MT in epithelial cells were comparable to those of TB both in the presence and absence of neutralizing antibodies (compare Figs 4D and 7B). This was consistent with the notion that both strains spread in epithelial cells predominantly via cell-to-cell spread over the first 12 days. Thus, while the high-levels of gH/gL/UL128-131 of ME clearly enhance the efficiency of the spread in epithelial cells, this does not appear to determine the mechanism, and these results suggest other intrastrain variable factors can affect cell-to-cell spread in this cell type.

### The RL13 glycoprotein tempers both cell-free and cell-to-cell spread

The RL13 ORF encodes an envelope glycoprotein that has been described as a selective, or preferential inhibitor of cell-free spread over cell-to-cell spread inasmuch as genotypes with inactivating mutations in RL13 arise during serial propagation of HCMV in both fibroblasts and epithelial cells, and this has been correlated with the appearance of more cell-free spread characteristics (40, 50, 55). While the ME recombinant used in the present studies harbors such an inactivating RL13 mutation, the RL13 expression status of the TB and TR BAC-clones was unclear. DNA sequencing confirmed that the RL13 ORF of both TB and TR was intact (data not shown), but lack of quality antibodies precluded confirmation of RL13 glycoprotein expression. Thus, to directly and fairly compare the effects of RL13 on the spread of all three strains, and to avoid potential selection of RL13 mutants in TB and TR during spread experiments, fibroblast and epithelial cell lines that express RL13 were engineered. Immunoblot analysis demonstrated that these cell lines expressed of both mature and immature proteoforms, and flow cytometry confirmed similar expression levels (Fig 8).

**Figure 8.**
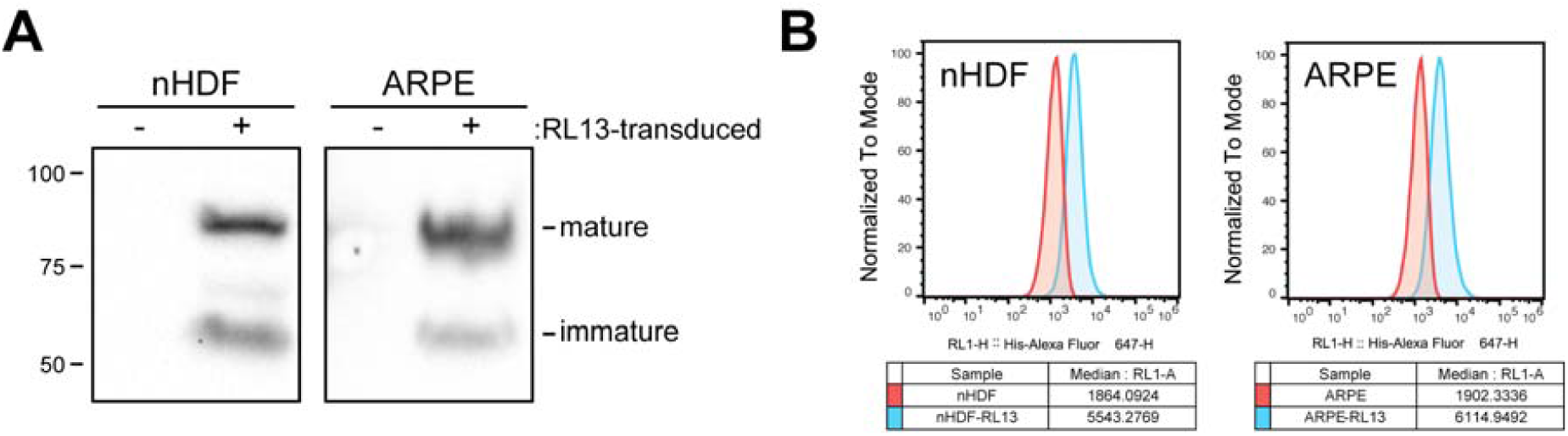
RL13-expressing fibroblasts and epithelial cell lines. nHDF fibroblasts and ARPE epithelial cells transduced with lentivectors encoding a C-terminal 6XHis tagged RL13 were analyzed by immunoblot (A) or flow cytometry (B) using anti-6XHis antibodies. Median fluorescence intensities for RL13-expressing nHDFs and ARPEs, and control cells are shown for comparison.

In fibroblasts, RL13 expression resulted in a comparable 40-50% reduction in spread rate for all three strains (Fig 9A). The relative effect of neutralizing antibodies on spread by TB or TR was unaltered by RL13 expression, but spread of ME was 2-fold more sensitive to neutralizing antibodies in the RL13-expressing fibroblasts. In epithelial cells, RL13 expression had no effect on the spread of TR but resulted in a modest 20% reduction for both TB and ME, although only for ME was this statistically reproducible (Fig 9B). Effects of RL13 expression on antibody sensitivity were also different in epithelial cells than in fibroblasts. RL13 expression increased the sensitivity of TB spread to antibody inhibition, but had little effect on TR or ME (Fig 9B). The notion that RL13 acts selectively or preferentially to restrict cell-free spread would have predicted that a greater fraction of the observed spread in RL13-expressing cells would be cell-to-cell and therefore less sensitive to neutralizing antibodies. On the contrary, results showed that sensitivity of spread to neutralizing antibodies was either unaffected or enhanced in RL13-expressing cells compared to the control cells. Thus, RL13 appears to exert its effect on HCMV spread in a manner that affects both the cell-free and the cell-to-cell mechanisms, not preferentially cell-free spread.

**Figure 9.**
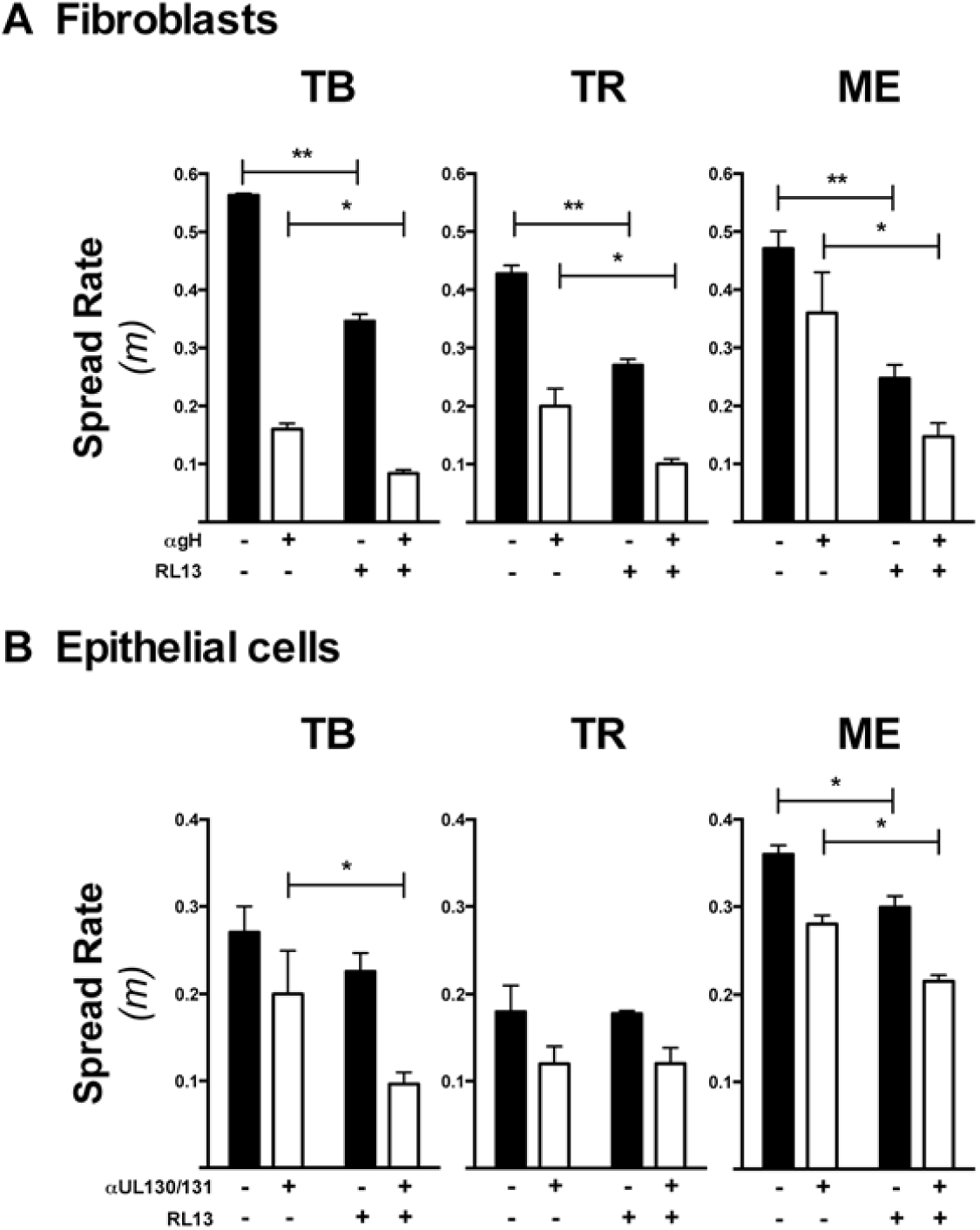
Effects of RL13 expression on spread of HCMV in fibroblasts and epithelial cells. Spread rates of TB, TR, or ME were determined as in Fig. 2 in RL13-expressing fibroblasts (A) or epithelial cells (B) and respective control cells in both the presence and absence of neutralizing antibodies anti-gH mAb 14-4b (A), or anti-UL130/131 rabbit sera (B). Plotted are average spread rates and standard deviations for three independent experiments. P-values were generated by unpaired two-tailed t-tests comparing RL13-expressing and control cells (*<0.05, **<0.01).

### RL13 exerts strain-dependent and cell type-dependent effects on the quantity and infectivity of progeny virus released to culture supernatants

Two plausible mechanisms by which RL13 expression could negatively affect the cell-free mode of spread include 1) reducing the numbers of cell-free progeny released to culture supernatants, and 2) reducing the infectivity of the cell-free progeny released. Both possibilities were tested using the RL13-expressing fibroblasts and epithelial cells. In fibroblasts, there were modest reductions in the numbers of progeny released to culture supernatants for TB and ME (Fig 10A). There were also modest reductions in detectable infectivity in culture supernatants of the RL-13 expressing cells (Fig 10B.) Taken together, the correlation between reduced numbers and reduced infectivity suggest that the specific infectivity of cell-free virions was not substantially affected by RL13-expression. For the efficient cell-free spreaders TB and TR, these reductions in cell-free progeny quantity and/or infectivity may help explain the reductions in spread rates described above. In contrast, the reduction of cell-to-cell spread for ME could not be attributed to reduced cell-free progeny quantity or infectivity.

**Figure 10.**
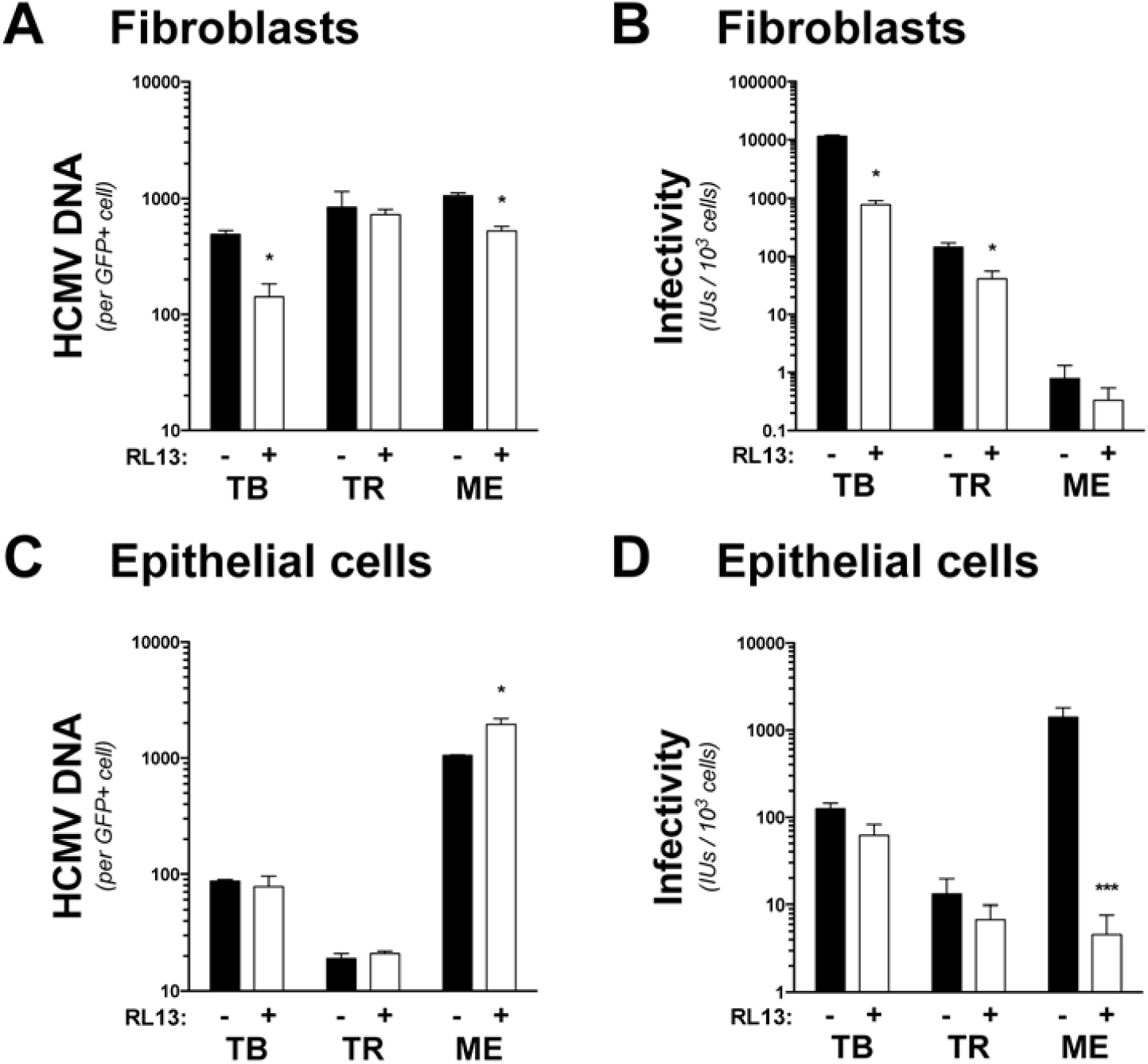
Quantity and infectivity of cell-free HCMV progeny virus from RL13-expressing fibroblasts and epithelial cells. RL13-expressing fibroblasts (A-B) and epithelial cells (C-D) or control cells were infected with HMCV strains TB, TR, or ME at MOI 1. At 7 days post infection the number of infected cells was determined by flow cytometry and culture supernatants were analyzed by qPCR for viral genomes (A and C) or by titration on matched producer cell type to quantify infectious units. Plotted are averages and standard deviation for three independent experiments. P-values were generated by unpaired two tailed t-tests comparing RL13-expressing and control cells (*<0.05, **<0.01, ***<0.001).

In epithelial cells, there were no effects on the numbers of supernatant progeny released for TB or TR, but there were actually more cell-free ME progeny produced from RL13-expressing cells (Fig. 10C). Despite the greater numbers of ME progeny in RL13-expressing epithelial cell supernatants, there were drastically lower levels of infectivity, indicating a substantial reduction in the per virion infectivity (Fig. 10D). However, as was noted above for fibroblasts, since the spread of ME in epithelial cells was determined to be predominately cell-to-cell, these effects on cell-free virus may not fully explain how RL13 tempers spread in this cell type.

The drastic reduction of ME cell-free infectivity due to RL13 expression in epithelial cells suggested effects on the envelope glycoproteins. In particular, gH/gL/UL128-131 is the only envelope glycoprotein currently known to be specifically required for efficient infection of epithelial cells. Thus, ME virions produced by RL13-expressing or control fibroblasts and epithelial cells were analyzed by immunoblot for the levels of gH/gL complexes. Consistent with our previous reports (Fig. 6B, and (51)), ME virions produced by fibroblasts had gH/gL predominately in the form of gH/gL/UL128-131 (Fig 11A). Strikingly, epithelial-derived ME contained substantially greater amounts of gH/gL/UL128-131 than fibroblast-derived ME (Fig 11A). In both cell types, RL13 expression resulted in a slight reduction in gH/gL/UL128-131, but the reductions were too small to provide a compelling explanation for the effects on virion infectivity observed. Levels of gB were also analyzed as a control, but surprising results were obtained. Antibodies used to detect gB were specific for both the 55 kDa portion of the furin-cleaved proteoform, and the the full length, uncleaved 165-170 kDa form and (56–58). In fibroblasts, ME virions contained almost entirely furin-cleaved gB, as evidenced by detection of only a 55kDa band (Fig 11B). This was not affected by RL13-expression. In contrast, epithelial-derived ME contained mostly uncleaved gB, and RL13-expression drastically reduced the amount of this proteoform (Fig 11B). Together, these results indicated that RL13 can affect the processing and incorporation of gH/gL complexes and gB into cell-free progeny, albeit in a cell type-dependent manner.

**Figure 11.**
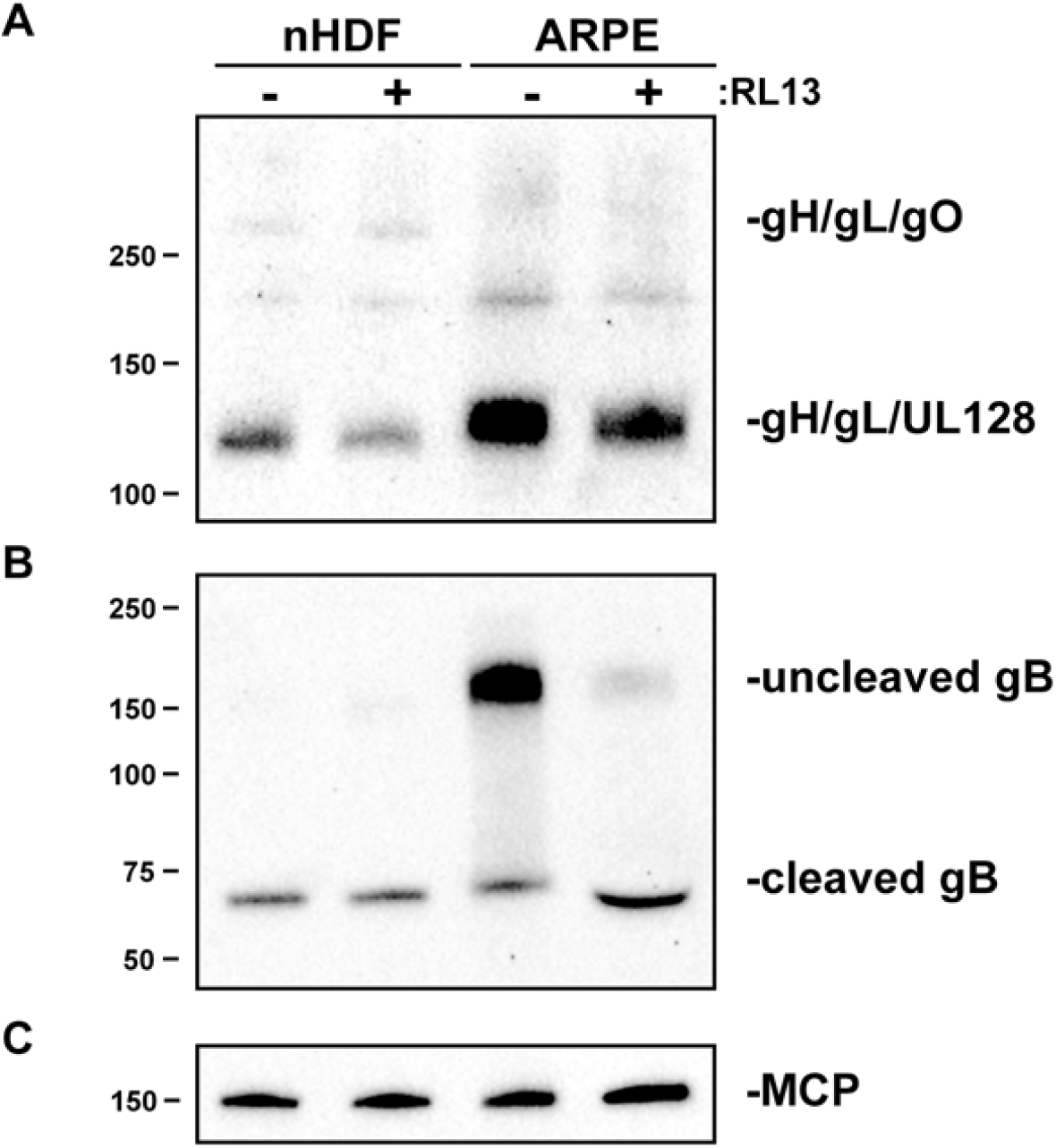
Effect of RL13 expression on levels of gH/gL complexes and gB in ME virions. Equal numbers (genomes/mL) of ME virions derived from either RL13-expressing or control fibroblasts or epithelial cells were separated by SDS-PAGE under non-reducing (A) or reducing (B and C) conditions and analyzed by immunoblot with antibodies directed against gL (A), gB (B), or major capsid protein (C).

## DISCUSSION

Distinctions between cell-free and cell-associated phenotypes of different HCMV strains and isolates have been noted, but the importance of different genetic loci involved in these spread modes remains to be understood. Sinzger et al. showed that during the initial rounds of cell-associated subculturing in fibroblasts, some clinical HCMV isolates formed small and tightly packed foci, but tended towards larger and more diffuse foci during later rounds of passage (59). The change in foci morphology coincided with the appearance of infectious virus in culture supernatants and the basic interpretation was that the adaptation of clinical HCMV isolates to the cell culture environment involved a shift from predominately cell-to-cell spread to cell-free spread. A more recent report by Galitska et al. noted considerable variation among clinical HCMV samples, with some displaying the characteristic small foci, but others showing larger diffuse foci even in early rounds of subculturing (60). This variation fits well with the growing evidence of genotypic diversity of HCMV populations within natural populations of HCMV (3, 4, 6, 8, 9) and with the earlier suggestion by Sinzger that the observed culture adaptation of HCMV clinical isolates might represent random sampling or purifying selection of preexisting genotypic variants, rather than the acquisition of *de novo* mutations (59). In line with this view, Subramanian et al. used a flow cytometry method to determine the increase in HCMV infected cells over time to better quantitate spread kinetics and found that some low passage clinical isolates spread more rapidly than others (61). They interpreted slow kinetics to represent a cell-associated-only phenotype and fast kinetics to represent a phenotype of both cell-associated and cell-free spread. Indeed, the faster spreading isolates were more sensitive to the presence of neutralizing antibodies. Together, these and other studies suggest considerable variation in the modes of spread exhibited by genetically distinct HCMV, but the exact genetic variations that determine these phenotypes remain unclear.

Here we applied a flow cytometry-based approach to compare the spread characteristics of three commonly studied and genetically distinct BAC clones of HCMV (TB, TR, and ME) in both fibroblast and epithelial cell cultures. It has been widely appreciated that TB and TR exhibit characteristics of cell-free spread, whereas ME is essentially restricted to cell-to-cell spread. Previous studies suggested that the distinction between these strains in the ability to spread cell-free in fibroblast cultures is due largely, if not entirely to the fact that cell-free TB and TR virions are log-folds more infectious than ME virions and that cell-free spread of ME in fibroblast cultures can be enhanced by transcriptional repression of epithelial/endothelial cell tropism factor gH/gL/UL128-131, which in turn leads to improved infectivity of cell-free ME virions (40) (38) (27). However, it has remained unclear whether the apparent cell-associated nature of ME is due simply to the poor cell-free infectivity, or reflects a specific ability to facilitate more efficient cell-to-cell spread than viruses like TB and TR that can spread efficiently cell-free. An important corollary consideration for our studies was that the UL128-131 and RL13 loci have been implicated as important for determining the cell-free or cell-associated nature of HCMV strains (24, 27, 40, 42) (40, 50, 55). The results presented here demonstrate that the cell-associated characteristic of ME is not simply due to the poor cell-free infectivity. Rather, ME is capable of a specifically efficient cell-to-cell mechanism of spread, TB is specifically inefficient at cell-to-cell spread, and TR may be of intermediate cell-to-cell spread capacity. Moreover, we demonstrate that cell-free and cell-associated spread characteristics are not determined by either the expression levels of UL128-131 or by RL13 expression, although these loci do influence the efficiency in strain-dependent and cell type-dependent manners. Thus, we suggest the involvement of other variable loci in distinct mechanisms of cell-free and cell-to-cell spread, and that in epistatic relationships among these loci govern the efficiency of these processes.

One candidate for a variable factor that might influence cell-free and cell-to-cell spread in an epistatic manner is gO, which is one of the more diverse proteins encoded by HCMV. There are 8 alleles of the UL74(gO) ORF that vary between 10-30% of predicted amino acids (62, 63) (64) (51). Mutational inactivation of UL74 in TR and TB dramatically reduced cell-free infectivity, but allowed cell-to-cell spread, albeit reduced (26) (65). In contrast, spread of a gO_null ME was indistinguishable from the parental (24). Together these observations suggested that gH/gL/gO is critical for cell-free spread, but cell-to-cell spread can be mediated by gH/gL/UL128-131 alone. However, it has more recently been reported that viruses lacking gH/gL/UL128-131 can spread cell-to-cell, but is dependent on the gH/gL/gO receptor, PDGFRa (43). Furthermore, a recent unpublished study by our group demonstrated that switching the UL74 allele in both TR and ME backgrounds could influence the efficiency of cell-to-cell spread, demonstrating that there were epistatic phenomena involved inasmuch as the influence of some UL74 alleles were observed in only one of the two genetic backgrounds (66).

Our finding that the level of gH/gL/UL128-131 expression did not affect the spread rate of ME in fibroblasts conflicts with a report by Stanton et al., which showed UL128-null ME plaques significantly larger than UL128-intact ME plaques (40). The apparent contradiction may be explained by the fact that our flow cytometry assays measured spread rates averaged over the first 12 days, as compared to the measurements of 21-day plaque sizes by Stanton et al. Linear regression analyses of the data collected by flow cytometry indicated that ME spread at a faster rate between days 3 and 6, and then began to slow after day 9, whereas the constant rate of cell-free spread demonstrated by TB and TR (Fig. 2). The slowdown displayed by ME may simply reflect a limiting number of potential uninfected target cells available via cell-to-cell spread during focal expansion. Alternatively, it could be that the cells at the periphery of an expanding ME foci rapidly become less permissive, for example via activation of the antiviral state. In any case, the increase in 21 day plaque size of a UL128-null ME is consistent with enhanced cell-free infectivity, but our analysis of spread rates over 12 days indicates this does not come at the expense of a distinct and comparably efficient mode of cell-to-cell spread. This interpretation is supported by the observation that the 12 day spread rate of ME was highly resistant to neutralizing antibodies regardless of gH/gL/UL128-131 expression level (Fig. 7) and is consistent with Laib Sampaio et al., who showed much greater cell-free dispersion of UL128-null ME foci, but more similar antibody resistant, cell-associated spread with or without UL128 (24).

In epithelial cells, repression of gH/gL/UL128-131 did reduce the 12 day spread rate for ME (Fig 7). This was consistent with reduced plaques sizes of UL128-null ME documented by Murrell et al., and likely reflects the important role of gH/gL/UL128-UL131 in epithelial cells (38). However, as observed in fibroblasts, spread in epithelial cells was still highly resistant to neutralizing antibodies when gH/gL/UL128-131 was repressed. This was different than Murrell et al., who concluded that spread of ME in epithelial cells was more sensitive to antibodies when gH/gL/UL128-131 was reduced (42). Again the discrepancy may indicate that analyses of 21-day plaque size likely accentuate the influence of cell-free infectivity and masks the contribution of cell-to-cell spread. Thus, by measuring average spread rates over 12 days, we find that gH/gL/UL128-131 levels are not the sole determinate of the cell-free versus cell-associated spread phenotype for ME. Rather, the increase in cell-free infectivity due to reduction in gH/gL/UL128-131 and the concomitant increase in gH/gL/gO, is independent of the highly efficient cell-to-cell mode of spread.

The RL13 glycoprotein was another candidate for a factor influencing the cell-free and cell-associated phenotypes. Previous reports have suggested that expression of RL13 limits HCMV replication in fibroblast, epithelial, and endothelial cell cultures through preferential effects on cell-free spread (24, 40, 50, 55). On the contrary, we find that RL13 tempers HCMV spread by either cell-free or cell-to-cell mechanisms and that this temperance was more pronounced in fibroblasts compared to epithelial cells (Fig. 9). Interestingly, RL13 expression dramatically reduced the infectivity of cell-free virions released by ME from epithelial cells, and this was consistent with the previous report showing low cell-free titers for RL13-intact ME in epithelial cell cultures compared to RL13 mutant ME (40). The physiological basis of this infectivity loss remains unclear, but it correlated with the observations that ME virions produced in RL13-expressing epithelial cells contained less full length, non-furin cleaved gB (Fig. 11). Strive et al. showed that furin cleavage was not required for the function of HCMV gB (67). Therefore, while the change in gB may not directly explain the change in infectivity, it is conceivable that RL13 could affect the processing and incorporation of other important envelope glycoproteins.

Our results may help to explain inconsistencies regarding the selection of UL128-131 and RL13 mutations during laboratory culturing of HCMV isolates, phenomenon that has been universally ascribed to HCMV, despite clear strain variability. While ME is highly sensitive to the selection of *de novo* UL128-131 in fibroblast cultures, TB and TR are remarkably stable in this respect (40, 50, 59). This disparity might simply reflect that many subculturing methods that including supernatant transfers, cell sonication, and high split-ratio transfer of intact infected cells, would be expected to heavily favor cell-free spread. Thus, given the highly efficient cell-to-cell spread of ME reported in this study, propagation by low split-ratio passage of intact, infected cells might reduce the selective bottleneck on loss of UL128-131 function to favor cell-free spread. Likewise, RL13 inactivating mutations have also been reported for many strains and clinical isolates in studies involving fibroblasts, epithelial, and endothelial cells (10, 55). The BAC clones of TB and TR have intact RL13 ORFs, but since expression of RL13 from transduced cells reduced replication by these strains, it may be that both strains carry mutations or polymorphisms in promoter or other regulatory sequences that impact RL13 expression. Moreover, our results suggest that the selection against RL13 may be more extensive in fibroblasts but less pronounced in epithelial cells.

Notably, a recent report showed that the presence of neutralizing antibodies during subculturing of clinical urine-derived HCMV in fibroblasts stabilized the UL128-131 and RL13 loci in the resultant virus populations (68). If these clinical isolates were similar to ME in their proclivity for cell-to-cell spread, it would seem logical that neutralizing antibodies would reduce the selective advantage of cell-free infectivity conferred by UL128-131 mutation. Indeed, it is interesting that ME was also isolated from a clinical urine sample (10). However, the preservation of RL13 seems less intuitive since RL13 impacts both cell-free and cell-to-cell modes of spread. Given the variable effects we observed of RL13 on different strains, these observations again seem to point to epistasis phenomena, where the relative effect of any given perturbation at one variable locus is influenced by variable physiology related to other polymorphic loci.

Our data also shed new light on the notion that the cell type might influence phenotypic properties in a non-genetic manner through changes to the protein composition of the progeny virus. Among the herpesviruses, these so called “producer cell effects” have been particularly well characterized for EBV, where the mechanism alternates the tropism of progeny virus between epithelial and B-cells (reviewed in (69)). Scrivano et al. suggested a similar phenomenon for HCMV based in part on data showing that progeny virus produced from endothelial cells had less UL128 protein in the envelope compared to fibroblast derived progeny (35). This is in stark contrast to our results that epithelial-derived HCMV contained greater amounts of gH/gL/UL128-131 than fibroblast-derived virus (Fig 9). There are several possible, non-mutually exclusive explanations for this discrepancy. This could reflect differences between strains as the prior study used TB, whereas our analyses used ME. There could also be differences between endothelial and epithelial cells. Finally, the Scrivano et al. analysis normalized virion immunoblots to gB, whereas we used major capsid protein (MCP). It is possible that incorporation of gB into progeny virus is affected by glycoprotein processing/secretory pathway differences between cell type, whereas plasticity in the number of subunits per capsid subunits seems less likely. Indeed, we observed differences in the proteolytic processing and virion incorporation of gB when ME was produced in fibroblast and epithelial cells.

In conclusion, we have characterized very different cell-free and cell-associated spread phenotypes among three commonly studied BAC clones of HCMV. Our data suggest that these phenotypic variances are manifest through the combined influenced of diversity at multiple genetic loci other than UL128-131 and RL13. How this phenotypic variation is represented *in vivo* remains unclear. There is little clear evidence to support the notion that HCMV is predominately cell-associated *in vivo*. The endemic nature, and the pleomorphic disseminated disease presentations suggest that HCMV is able to thrive in many different bodily environments as it spreads within and among individual human hosts. Copious amounts of cell-free virus are shed in urine, saliva, and breast milk, and this is likely a major route of transmission between individuals, whereas HCMV in the blood is likely highly associated with leukocytes (70–72). Thus, it seems likely that both cell-free and cell-associated modes of spread are important to the natural history of HCMV.

## MATERIALS AND METHODS

### Cell lines

Primary neonatal human dermal fibroblasts (nHDF; Thermo Fisher Scientific), RL13-nHDFs (nHDFs transduced with lentiviral vectors encoding RL13 of HCMV strain Merlin, selected with puromycin resistance), nHDF-tet (nHDFs transduced with retroviral vectors encoding the tetracycline repressor protein, selected with puromycin resistance, MRC-5 fibroblasts (American Type Culture Collection; CCL-171), and HFFF-tet (40) (provided by Richard Stanton, Cardiff University, United Kingdom) were grown in Dulbecco’s modified Eagle’s medium (DMEM)(Sigma) supplemented with 6% heat-inactivated fetal bovine serum (FBS) (Rocky Mountain Biologicals, Missoula, MT, USA) and 6% Fetalgro® (Rocky Mountain Biologicals, Missoula, MT, USA). Retinal pigment epithelial cells (ARPE19)(American Type Culture Collection, Manassas, VA, USA), ARPE-tet (ARPE19 cells transduced with retroviral vectors encoding the tetracycline repressor protein, and selected by puromycin resistance), and RL13-ARPE (transduced with lentiviral vectors encoding RL13 of HCMV strain Merlin, selected with puromycin resistance) were grown in a 1:1 mixture of DMEM and Ham’s F-12 medium (DMEM:F-12)(Gibco) and supplemented with 10% FBS.

### Retroviral vectors

The Tet repressor protein bearing a nuclear localization signal was extracted by PCR from the integrated sequence in the HFFF-tet chromosomal DNA and cloned into the same pMXs retrovirus vector used to construct HFFF-tet (40). The pMXs retrovirus vector plasmid was a gift from Dr. Toshio Kitamura at the Institute of Medical Science, University of Tokyo (73). The tet-containing vector plasmid was transformed in 293T cells together with the pUMVC and pMD2.G helper plasmids. The pUMVC helper plasmid was a gift from Bob Weinberg (Addgene plasmid # 8449) (74). Two days after transformation, the retroviral particles in the supernatant were purified from cell debris through syringe filtration and centrifugation. After titration, the particles were used to transduce low passage nHDF or ARPE-19 cells. After a week of puromycin selection, cells aliquots were tested for tetR expression after transfection of a firefly luciferase tetR-reporter system and aliquots were stored in liquid nitrogen until further usage. The codon-optimized RL13 with an intact ORF from Merlin HCMV strain (NCBI ref YP_081461) was constructed by Gibson Assembly and used to replace the eGFP ORF in the pLJM1-EGFP lentiviral transfer vector plasmid. The pLJM1-EGFP plasmid was a gift from David Sabatini (Addgene plasmid # 19319) (75). The RL13-containing vector plasmid was transformed in 293T cells together with three lentiviral helper plasmids. The pMDLg/pRRE, pRSV-Rev, and pMD2.G helper plasmids were a gift from Didier Trono (Addgene plasmids # 12251, # 12253, # 12259, respectively) (76). Two days after transformation, the lentiviral particles in the supernatant were purified from cell debris thru syringe filtration and centrifugation. After titration, the particles were used to transduce either low passage nHDF or ARPE-19 cells. After a week of puromycin selection, cells were tested for RL13 expression and aliquots were stored in liquid nitrogen until further usage.

### HCMV

All human cytomegalovirus (HCMV) strains were derived from bacterial artificial chromosome (BAC) clones. BAC clone TB40/e (BAC4) was provided by Christian Sinzger (University of Ulm, Germany) (41). BAC clone TR was provided by Jay Nelson (Oregon Health and Sciences University, Portland, OR, USA) (77). BAC clone Merlin (ME)(pAL1393), which contains tetracycline operator sequences within the transcriptional promoter of UL130 and UL131, was provided by Richard Stanton (Cardiff University, Cardiff, United Kingdom) (40). All BAC clones were modified to express green fluorescent protein (GFP) with En passant recombineering (78) by replacing US11 with the eGFP gene. The constitutive expression of eGFP allows the monitoring of HCMV infection early and is strain-independent. Infectious HCMV was recovered by electroporation of BAC-DNA into HFFFs, as previously described (26). For infectious unit (IU) determination, viruses were serial diluted and infectivity was determined on fibroblasts or epithelial cells using flow cytometry 48 hours post infection.

### Antibodies

Monoclonal antibodies specific to HCMV major capsid protein (MCP), gH (14-4b), and gB (27-156) were provided by Bill Britt (University of Alabama, Birmingham, AL) (52, 57, 79). 14-4b was purified by FPLC prior to use. Rabbit polyclonal sera against HCMV gL, UL130, and UL131 were provided by David Johnson (Oregon Health and Sciences University, Portland, OR) (17).

### Vrial spread assays

Approx. 1×10^5^ (3×10^5^) nHDFs (ARPEs) were seeded into 6-well culture plates and allowed to grow to confluence. Confluent monolayers of nHDFs or ARPE cells were inoculated with 100-1000 IUs of strains TB, TR, or ME at for 4hrs at 37**°**C. Cells were then washed extensively with PBS and cultured in the appropriate growth medium supplemented with 2% FBS. Viral spread in the presence or absence of neutralizing antibodies or ganciclovir was monitored over 12 days by flow cytometry. For fibroblasts, 100ug/mL anti-gH 14-4b was used; for epithelial cells, a 1:1000 dilution of anti-UL130/131 rabit sera. 50uM ganciclovir was used on both cell types. All experiments were performed in triplicate and a minimum of 3 experiments was conducted for each condition. Spread rates were determined by plotting LN GFP+ cells over time and fitting the data to the log-linear rate expression LN(I)_t_ = *m*(t) + LN(I)_0_ where (I) is the number of GFP+ cells, (t) is the time in days, and *m* is the spread rate.

### Flow cytometry

Recombinant GFP-expressing HCMV-infected cells were washed twice with PBS and lifted with trypsin. Trypsin was quenched with DMEM containing 10% FBS and cells were spun at 500xg for 5 min at RT. Cells were fixed in PBS containing 2% paraformaldehyde for 10 min at RT, then washed and resuspended in PBS. Samples were analyzed using an AttuneNxT flow cytometer. Cells were identified using FSC-A and SSC-A, and single cells were gated using FSC-W and FSC-H. BL-1 laser (488nm) was used to identify GFP+ cells, and only cells with median GFP intensities 10-fold above background were considered positive. RL13 expression was measured using an intracellular staining kit (Thermo) and an anti-6x His antibody conjugated to AlexaFluor-647 (Thermo) using the RL-2 laser (647nm).

### qPCR

The real-time quantitative PCR (qPCR) assay used to quantify viral or cellular DNA molecules was performed as previously described (27). Briefly, HCMV DNA inside cell-free particles was purified using the PureLink Viral RNA/DNA mini kit (Thermo Scientific). A region within UL83 conserved among ME, TR, and TB was chosen as the HCMV-specific amplicon, and viral genomes were quantified by SYBR green qPCR as previously described. Standard curves were performed using serial dilutions of a single PCR DNA band containing the sequences of the viral UL83 and cellular beta2-microglobulin (see below) amplicons and the corresponding set of primers. Finally, the concentration of HCMV DNA genomes in the supernatant was extrapolated from the UL83 standard curve and expressed as genome molecules per ml of media (genome/ml). Total intracellular or cytoplasmic HCMV DNA was quantified with the UL83-based qPCR after extracting viral and cellular DNA from infected cells with the PureLink Genomic DNA mini kit (Thermo Scientific). We also measured the chromosomal DNA molecules present in these samples with an amplicon located in the human beta-2 microglobulin gene (80). These chromosomes numbers were used to correct for the number of cells when measuring the whole cell, nuclear, and cytoplasmic genomes from different samples. To obtain these fractions, infected cells were subjected to a Subcellular Fractionation Kit for cultured cells (Thermo Scientific) to purify the cytoplasmic HCMV DNA from the excess of nuclear HCMV DNA. Lastly, genomes present in the cytoplasmic fraction were purified using the Genomic DNA mini kit and their number was measured with the UL83-based qPCR assay.

### Particle production

Production of intracellular and extracellular HCMV genomes was determined over a time course of 4 days. Briefly, nHDFs and ARPEs were infected with HCMV strains TB, TR, or ME at an MOI of 1, and the cells were extensively washed 3 days post infection. At 3, 5, and 7 dpi culture supernatants were collected and spun at 500xg for 10 minutes at RT, while cells were lifted with trypsin and then quenched with culture media containing 10% FBS. Cells were spun at 500xg and resuspended into aliquots. Cells were either resuspended in PBS for whole cell analysis, or fractionated for nuclear and cytoplasmic analysis using the Subcellular Fractionation Kit for Cultured Cells (Thermo). Cells were also sonicated to release cell-associated virus for infectivity analysis, or processed for flow cytometry to determine the number of GFP+ cells. The number of HMCV genomes in the whole cell, nuclear, cytoplasmic, and supernatant fractions was measured by qPCR, normalized to load, and divided by the number of GFP+ cells. The total infectivity of supernatant and sonicated fractions was determined by flow cytometry 48 hours after inoculation of the producer cell type. The accumulation at day 7 from nHDFs and ARPEs is shown in Fig 5.

### Immunoblot analyses

Cell-free virions were solubilized in a buffer containing 20mM Tris-buffered saline (TBS) (pH 6.8) and 2% SDS. Protein samples were separated by SDS-polyacrylamide gel electrophoreses (SDS-PAGE) and electrophoretically transferred to polyvinylidene difluoride membranes in a buffer containing 10mM NaHCO_3_ and 3mM Na_2_CO_3_ (pH 9.9) and 10% methanol. Transferred proteins were first probed with MAbs or rabbit sera, then anti-mouse or anti-rabbit secondary antibodies conjugated to horseradish peroxidase (Sigma), and Pierce ECL-Western Blotting substrate (Sigma). Chemiluminescence was detected using a Bio-Rad ChemiDoc MP imaging system.

## ACKNOWLEDGEMENTS

We are grateful to Richard Stanton, Bill Britt, Jay Nelson, and David Johnson, for generously supplying HCMV BAC clones, antibodies, and cell lines as indicated in the Materials and Methods. Additionally, we are grateful to the Center for Biomolecular Structure and Dynamics (CBSD), University of Montana, Missoula, MT, for purification of monoclonal antibodies, and well as the Flow Cytometry Core of the Center for Environmental Heath Sciences (CEHS), University of Montana, Missoula, MT, for guidance on experimental design, acquisition, and analysis of the flow cytometry-based approaches used for this study.

This work was supported by grant from the National Institutes of Health to B.J.R (R01AI097274), a fellowship from American Heart Association to E.P.S (17POST33350043) and a National Institutes of Health CoBRE award to Center for Biomolecular Structure and Dynamics at University of Montana (PG20GM103546).

Experiments were designed by E.P.S, B.J.R, and J.L., and performed by E.P.S, J.L, L.Z.D., C.P., J.P, and Q.Y., and the manuscript was prepared by B.J.R., E.P.S., and J.L.

## FIGURE LEGENDS

**Figure 1. Comparison of focal spread patterns among distinct HCMV strains.** Confluent monolayers of fibroblasts (A) or epithelial cells (B) were infected at MOI 0.001 with GFP-expressing HCMV TB, TR, or ME. Foci were documented by fluorescence microscopy at 6, 12, and 18 days post infection. Four representative fields (10X) are shown for each.

**Figure 2. Quantitation of HCMV spread by in fibroblasts and epithelial cells.** Confluent 9.5cm^3^ monolayers of fibroblasts (A) or epithelial cells (D) were infected at MOI 0.001 with GFP-expressing HCMV strains TB, TR, or ME. The number of infected cells at 3, 6, 9, and 12 days post infection was determined by flow cytometry. (B and E) LN GFP+ cells is plotted at each time point shown in panels A and D, respectively. (C and F) The average spread rate (LN GFP+ cells per day) as the slope (m) of the regression lines in panels B and E, respectively. All experiments were performed at least three times and error bars represent standard deviation between all experiments (omitted if smaller than the data marker), and p-values were generated using unpaired, two-tailed t-tests (*<0.05, **<0.01, ***<0.001).

**Figure 3. Antibody neutralization of cell-free HCMV.** Equal numbers (genomes/mL) of fibroblast-derived (A) or epithelial-derived (B) HCMV TB, TR, or ME virions were incubated with multiple concentrations of anti-gH mAb 14-4b (A) or anti-UL130/131 rabbit sera (B) for 1hr at RT. Remaining infectivity was determined by titration on the matched producer cell type, and plotted as percent of no antibody control. All experiments were performed in triplicate and error bars represent standard deviation.

**Figure 4. Effects of neutralizing antibodies on spread of HCMV strains in fibroblasts and epithelial cells.** Confluent 9.5cm^3^ monolayers of fibroblasts or epithelial cells were infected at MOI 0.001 with GFP-expressing HCMV strains TB, TR, or ME and allowed to spread in the presence of either anti-gH mAb 14-4b (A-C), anti-UL130/131 rabbit sera (D-F), or ganciclovir (all panels). The average spread rates over 12 days (LN GFP+ cells per day; as in Fig 2) for TB, TR, and ME on fibroblasts (A-C) and epithelial cells (D-F) are shown. Conditions where no spread was detected are designated “n.d.”. All experiments were performed three times and error bars represent standard deviation. P-values were generated by unpaired, two-tailed t-test vs. mock (*<0.05, **<0.01, ***<0.001).

**Fig 5. Production and infectivity of cell-free and cell-associated HCMV progeny virus.** Replicate cultures of fibroblasts or epithelial cells were infected at MOI 1 with TB, TR, or ME. 7 days later infected cells were quantified by flow cytometry for GFP expression. *(A and C).* Cultures were fractionated into whole cell (WC), nuclear (N), cytoplasmic (C), and supernatant (S), and viral genomes were determined by qPCR. The number of HCMV genomes in each fraction is plotted per GFP+ cell. *(B and D).* Infectious units were determined from whole cell sonicates (WC) and culture supernatants (S) by titration on the matched producer cell type. Total infectious units are plotted per 1000 GFP+ cells. *All panels.* Error bars represent the standard deviation of three separate experiments, and p-values were generated using unpaired, two-tailed t-tests (*<0.05, **<0.01, ***<0.001, ****<0.0001).

**Fig 6. Repression of gH/gL/UL128-131 expression from HCMV ME in fibroblasts and epithelial cells**. (A) Tetracycline-repressor protein (tetR) expressing fibroblasts (HFFFtet, nHDFtet) or epithelial cells (ARPEtet) were transfected with TetO-firefly luciferase reporter and the relative luciferase activity was measured. HFFFtet were treated with doxycycline (+) or left untreated (-). (B) Non-reducing (top) or reducing (bottom) immunoblot analyses of MT or ME virions derived from TetR-expressing or control cells, respectively. Blots were probed with anti-gL antibodies (top) or anti-MCP antibodies (bottom). (C) Specific infectivity (genomes/IU) of MT and ME virions derived from culture supernatants of TetR-expressing or control cells, respectively. Titrations of fibroblasts-derived virus were performed of nHDFs, epithelial-derived virus on ARPEs. All experiments were performed in triplicate and error bars represent standard deviation. P-values were generated by unpaired, two-tailed t-test (*<0.05, **<0.01, ***<0.001).

**Figure 7. Effect of gH/gL/UL128-131 repression on HCMV ME spread in fibroblasts and epithelial cells**. Confluent 9.5cm^3^ monolayers of (A) nHDF or nHDFtet, for ME and MT, respectively, (B) HFFFtet (+) or (-) doxycycline, for ME and MT, respectively, or (C) ARPE or ARPEtet, for ME and MT, respectively were infected with at MOI 0.001 with GFP-expressing HCMV ME/MT. Spread rates in the presence of either anti-gH mAb 14-4b (A and B), or anti-UL130/131 rabbit sera (C) were determined as described in Figure 2. Plotted are averages and standard deviaitons for three independent experiments. P-values were generated by unpaired, two-tailed t-test (*<0.05, **<0.01).

**Figure 8. RL13-expressing fibroblasts and epithelial cell lines.** nHDF fibroblasts and ARPE epithelial cells transduced with lentivectors encoding a C-terminal 6XHis tagged RL13 were analyzed by immunoblot (A) or flow cytometry (B) using anti-6XHis antibodies. Median fluorescence intensities for RL13-expressing nHDFs and ARPEs, and control cells are shown for comparison.

**Figure 9. Effects of RL13 expression on spread of HCMV in fibroblasts and epithelial cells.** Spread rates of TB, TR, or ME were determined as in Fig. 2 in RL13-expressing fibroblasts (A) or epithelial cells (B) and respective control cells in both the presence and absence of neutralizing antibodies anti-gH mAb 14-4b (A), or anti-UL130/131 rabbit sera (B). Plotted are average spread rates and standard deviations for three independent experiments. P-values were generated by unpaired two-tailed t-tests comparing RL13-expressing and control cells (*<0.05, **<0.01).

**Figure 10. Quantity and infectivity of cell-free HCMV progeny virus from RL13-expressing fibroblasts and epithelial cells.** RL13-expressing fibroblasts (A-B) and epithelial cells (C-D) or control cells were infected with HMCV strains TB, TR, or ME at MOI 1. At 7 days post infection the number of infected cells was determined by flow cytometry and culture supernatants were analyzed by qPCR for viral genomes (A and C) or by titration on matched producer cell type to quantify infectious units. Plotted are averages and standard deviations for three independent experiments. P-values were generated by unpaired two tailed t-tests comparing RL13-expressing and control cells (*<0.05, **<0.01, ***<0.001).

**Figure 11. Effect of RL13 expression on levels of gH/gL complexes and gB in ME virions.** Equal numbers (genomes/mL) of ME virions derived from either RL13-expressing or control fibroblasts or epithelial cells were separated by SDS-PAGE under non-reducing (A) or reducing (B and C) conditions and analyzed by immunoblot with antibodies directed against gL (A), gB (B), or major capsid protein (C).

